# The outer membrane and peptidoglycan layer form a single mechanical device balancing turgor

**DOI:** 10.1101/2023.04.29.538579

**Authors:** Michaël Deghelt, Seung-Hyun Cho, Sander K. Govers, Arne Janssens, Alix Dachsbeck, Han K. Remaut, Jean-François Collet

## Abstract

Bacteria are subject to a substantial concentration differential of osmolytes between the interior and exterior of the cell, which results in cytoplasmic turgor pressure. Failure to mechanically balance turgor pressure causes cells to burst. Here, we show that in Gram-negative bacteria, the outer membrane and peptidoglycan layer function together to resist turgor: when attached to each other, these two layers form a robust mechanical unit that allows pressure build-up in the periplasmic compartment, which in turn balances cytoplasmic turgor across the inner membrane, preventing cell death. Thus, the peptidoglycan layer is necessary but not sufficient to maintain turgor, which challenges the general view that protecting cells from bursting is the specific task of the peptidoglycan cell wall.

**ONE-SENTENCE SUMMARY:** The peptidoglycan and outer membrane are interconnected layers that cooperate to balance cytoplasmic turgor.

## INTRODUCTION

The cell envelope of Gram-negative (diderm) bacteria consists of two membranes, the cytoplasmic and outer membranes, which delimit a space known as the periplasm (*1*). A thin layer made of peptidoglycan, a polymer consisting of linear chains of glycans interconnected by short peptides (*2*), is present in the periplasm, embedded within the two membranes. All three envelope layers are necessary for growth and survival. The peptidoglycan layer, often referred to as the cell wall, contributes mechanical strength to cells, enabling them to resist osmotic challenges: when exposed to antibiotics that block the assembly of the peptidoglycan sacculus, cells cannot withstand the difference in osmotic pressure with the environment and lyse (*3*). The outer membrane, an asymmetric membrane with lipopolysaccharides in the outer leaflet and phospholipids in the inner leaflet (*4*), is primarily recognized for its role as an efficient permeability barrier that prevents the entry of toxic molecules into the cell (*5*). However, recent work has shown that the outer membrane also contributes to the mechanical properties of the cell envelope, functioning as a load-bearing structure important for cell stiffness (*6*). In *Escherichia coli*, as in several other bacterial species (*7*–*10*), the outer membrane and the peptidoglycan layer are interconnected: covalent and non-covalent interactions between outer membrane proteins and the peptidoglycan tether the membrane to the sacculus. These connections are important for maintaining outer membrane integrity (*11, 12*) and preventing the formation of outer membrane vesicles (*13*).

Although the last years have seen major progress in our understanding of how bacteria build and maintain the outer membrane and peptidoglycan layer, crucial problems remain unsolved. For instance, despite recent molecular insights (*14, 15*), we still have only a poor understanding of how the pathways that assemble the two outermost envelope layers are coordinated at the spatiotemporal level. Moreover, the manner in which the various outer membrane–peptidoglycan connectors impact cell physiology remains largely unexplored. Finally, although numerous well-established results have emphasized the crucial function of the outer membrane, the reasons for its essentiality are still unclear (*16*). Intriguingly, the only machineries that are essential in the outer membrane are those required for its own assembly (*16*). Here, we report an essential role of the outer membrane: when attached to the peptidoglycan layer, the outer membrane serves as part of a mechanical device that protects cellular integrity during osmotic challenges, preventing cell lysis. Our study overturns the widely accepted view that only the peptidoglycan layer mechanically maintains turgor pressure and suggests a model in which build-up of periplasmic pressure is crucial to balance cytoplasmic turgor and enable cell survival.

## RESULTS

### Role of the outer membrane–peptidoglycan connection in osmoprotection

At the beginning of this research, we found that cells lacking both the lipoprotein Lpp and the β-barrel OmpA, two major outer membrane–peptidoglycan connecters, exhibit a severe survival defect when exposed to hypoosmotic shocks (Fig. 1A–B). The phenotype of the single *lpp* and *ompA* deletion mutants was similar to that of the wild type (Fig. 1B). Expression at physiological levels of Lpp and OmpA variants with point mutations (Lpp_ΔK58_ and OmpA_R256E_) (*7, 17*) preventing peptidoglycan attachment did not restore the wild-type phenotype (Fig. 1B, fig. S1), indicating that the sensitivity to osmotic down-shifts specifically results from a lack of outer membrane–peptidoglycan connection and not from changes in the content of the membrane proteome. These results were unexpected because the current model of osmoresistance proposes that the cytoplasmic membrane expansion due to water influx under hypoosmotic shock is constrained by the peptidoglycan cell wall (*18*). Further, previous research has shown that mechanosensitive channels (MSCs) in the cytoplasmic membrane open when membrane tension increases, releasing ions and hydrated solutes to the periplasm, which limits the increase in cytoplasmic turgor pressure (*19*). Thus, our current understanding does not explain why connecting the outer membrane to the peptidoglycan layer is crucial for osmoprotection.

**Fig. 1.**
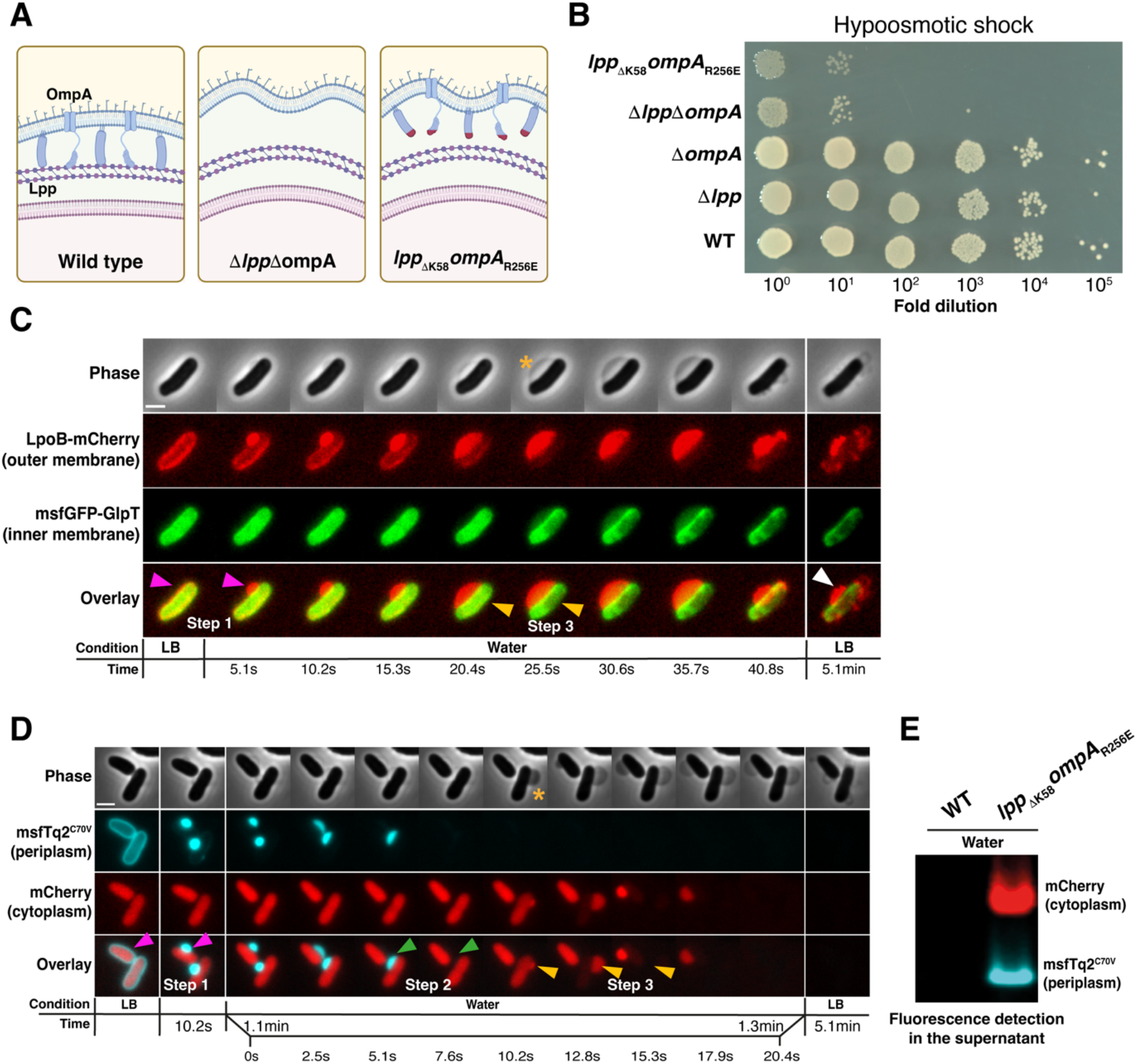
Connecting the outer membrane to the peptidoglycan layer is crucial for osmoprotection. (**A**) Lpp and OmpA attach the outer membrane to the peptidoglycan layer. In cells in which *lpp* and *ompA* are deleted or mutated (*lpp*_ΔK58_*ompA*_R256E_), attachment of the outer membrane to the peptidoglycan layer is prevented. (**B**) Whereas wild-type (WT) and single mutant cells survive a hypoosmotic shock (from LB to water), Δ*lpp*Δ*ompA* and *lpp*_ΔK58_*ompA*_R256E_ mutants exhibit a severe survival defect. Cells grown to exponential phase in LB were washed in water, serially diluted as indicated, and spotted on an LB plate. (**C**) The impact of hypoosmotic shock on cells with an impaired outer membrane–peptidoglycan connection was analyzed using microfluidics. *lpp*_ΔK58_*ompA*_R256E_ cells expressing LpoB-mCherry (outer membrane) and msfGFP-GlpT (inner membrane) were shifted from LB to water and remained in water for 5 min before being shifted back to LB. Snapshots were taken at the indicated times. Exposure to water causes rapid bulge formation (step 1, <5.1 s, purple arrows); the bulge increases in size over time. The msfGFP-GlpT signal is perturbed at 25.5 s (step 3, orange arrows), when the phase contrast signal of the periplasm becomes denser (yellow star). When shifted back to LB, LpoB-mCherry relocalizes around the cell body (white arrow), but most cells (97.9%±3.8%; n=2098 cells) do not undergo plasmolysis, indicating cell death. Scale bar: 2 μm. Images were acquired every 2.55 s. (**D**) The impact of hypoosmotic shocks on cells with an impaired outer membrane–peptidoglycan connection was analyzed using microfluidics. *lpp*_ΔK58_*ompA*_R256E_ cells expressing msfTq2^C70V^ (periplasm) and mCherry (cytoplasm) were shifted from LB to water and remained in water for 5 min before being shifted back to LB. Snapshots were taken at the indicated times. Hypoosmotic shock causes msfTq2^C70V^ to relocalize to the nascent bulge (step 1, purple arrows); the msfTq2^C70V^ signal is then lost (step 2, green arrows) (mean time=13.6 s after water shock; n=1747 cells). Approximately 30 s after loss of the msfTq2^C70V^ signal (mean time=30.1 s, n=1977 cells), the mCherry signal rapidly moves to the periplasm before being released out of the cell (step 3, orange arrows). The phase contrast signal of the periplasm becomes denser when mCherry moves to the periplasm; it decreases upon mCherry release (orange star). When shifted back to LB, most cells (99.8%±0.1%; n=2858 cells) do not respond to the osmotic upshift, indicating cell death. Scale bar: 2 μm. Images were taken every 2.55 s. (**E**) Periplasmic msfTq2^C70V^ and cytoplasmic mCherry are released in the supernatant when *lpp*_ΔK58_*ompA*_R256E_ but not wild-type (WT) cells are exposed to a hypoosmotic down-shift. Cell supernatants after hypoosmotic shock were loaded onto a native gel and imaged by fluorescence.

To obtain insights into this intriguing observation, we used a microfluidics-based assay to investigate the importance of connecting the outer membrane to the peptidoglycan layer under osmotic downshifts at the single-cell level. In one set of experiments, we labeled the cytoplasm and periplasm of wild-type and mutant *lpp*_βK58_*ompA*_R256E_ cells with genetically encoded mCherry and msfTq2^C70V^, respectively. In another set of experiments, we labeled the cytoplasmic membrane by expressing msfGFP fused to the glycerol-3-phosphate transporter GlpT (*20*) and used the outer-membrane lipoprotein LpoB fused to mCherry to monitor the outer membrane (*21, 22*). When the wild-type was shifted from lysogeny broth (LB) medium to water (hypoosmotic shock), the four fluorescent markers were unchanged (fig. S2, movies S1–2). When the wild-type shifted back to LB, plasmolysis occurred, indicating that wild-type cells remain viable when exposed to an osmotic up-shift (*23*), in agreement with the results in Fig. 1B. The behavior of the *lpp*_βK58_*ompA*_R256E_ mutant upon water shock was strikingly different (Fig. 1C–D, fig. S3, movies S3–4). First, following water influx, the outer membrane, marked by LpoB-mCherry, formed a large bulge in which periplasmic msfTq2^C70V^ accumulated (step 1); msfTq2^C70V^ was then released to the outside (after ∼13.6 s; step 2). Shortly afterward (∼30.1 s; step 3), cytoplasmic mCherry was transferred to the periplasm and, from there, out of the cell. The cytoplasmic membrane was also impacted in step 3, as revealed by perturbations in the distribution of msfGFP-GlpT (Fig. 1C, fig. S4). We observed that the phase contrast signal of the periplasm increased concomitantly with the transfer of mCherry, suggesting the passage of large amounts of cytoplasmic components to this compartment (Fig. 1C–D, fig. S4, movies S3–4). However, this increase was transient, hinting to the subsequent release of content to the media.

Accordingly, we detected mCherry, msfTq2^C70V^, and a large number of cytoplasmic and periplasmic proteins in the supernatant of the *lpp*_βK58_*ompA*_R256E_ mutant following water shock (Fig. 1E, fig. S5). When shifted back to the original osmolarity, the mutant cells did not undergo plasmolysis or resume growth, indicating that they were dead. While the death of *lpp*_βK58_*ompA*_R256E_ cells from the osmotic down-shift as observed by microfluidics is consistent with the results in Fig. 1B, the preservation of cell shape and density is notably different from the cell burst induced by β-lactams (*3*). To gain further insights into the effects of water shock on the cell envelope, we employed cryo-electron microscopy (cryo-EM). Our analysis confirmed the maintenance of cytoplasmic density and a uniform peptidoglycan layer, while also revealing that the outer membrane becomes significantly detached from the cell body when *lpp*_βK58_*ompA*_R256E_ cells, but not wild-type cells, are subjected to osmotic down-shifts (fig. S6). However, insights into the changes occurring in the cytoplasmic membrane and impacting msfGFP-GlpT distribution could not be obtained.

### Outer membrane–peptidoglycan model for maintaining osmoprotection

We interpret the results presented in Fig. 1 as follows (Fig. 2, top). Upon an osmotic down-shift, rapid water influx into the cytoplasm increases turgor, which applies an outward pressure on the cytoplasmic membrane (*24*). The resulting increase in membrane tension activates the MSCs (*19*), allowing the passage of osmotically active solutes to the periplasm. Solutes smaller than ∼600 Da are able to freely diffuse across the outer membrane (*25*). However, previous work has shown that such solutes accumulate, at least transiently, in the periplasm to balance the charged functional groups present on the macromolecules (proteins, phospholipids, peptidoglycans, and oligosaccharides such as osmoregulated periplasmic glucans (*26*)) trapped in this compartment. The uneven distribution of small solutes across a semi-permeable membrane causes the Donnan effect (*18, 27*). In turn, this accumulation of small solutes attracts more water into the periplasm. In wild-type cells, the outer membrane is attached to the peptidoglycan layer, which limits the expansion of the periplasmic volume and allows periplasmic turgor to build up. An inward pressure is then applied on the cytoplasmic membrane, balancing cytoplasmic turgor and perhaps promoting the closure of MSCs (Fig. 2, top). These cells resist hypoosmotic shocks. In contrast, when the outer membrane is not attached to the peptidoglycan layer, the influx of water in the periplasm due to increased concentrations of active solutes pushes the outer membrane apart from the cell body, which leads to an increase in periplasmic volume through the formation of a large bulge from which the soluble periplasmic content exits the cell, preventing the build-up of periplasmic turgor (Fig. 2, bottom). As a result, no inward pressure is applied on the cytoplasmic membrane, which becomes destabilized and releases cytosolic components to the periplasm; in such cases, the cells die.

**Fig. 2.**
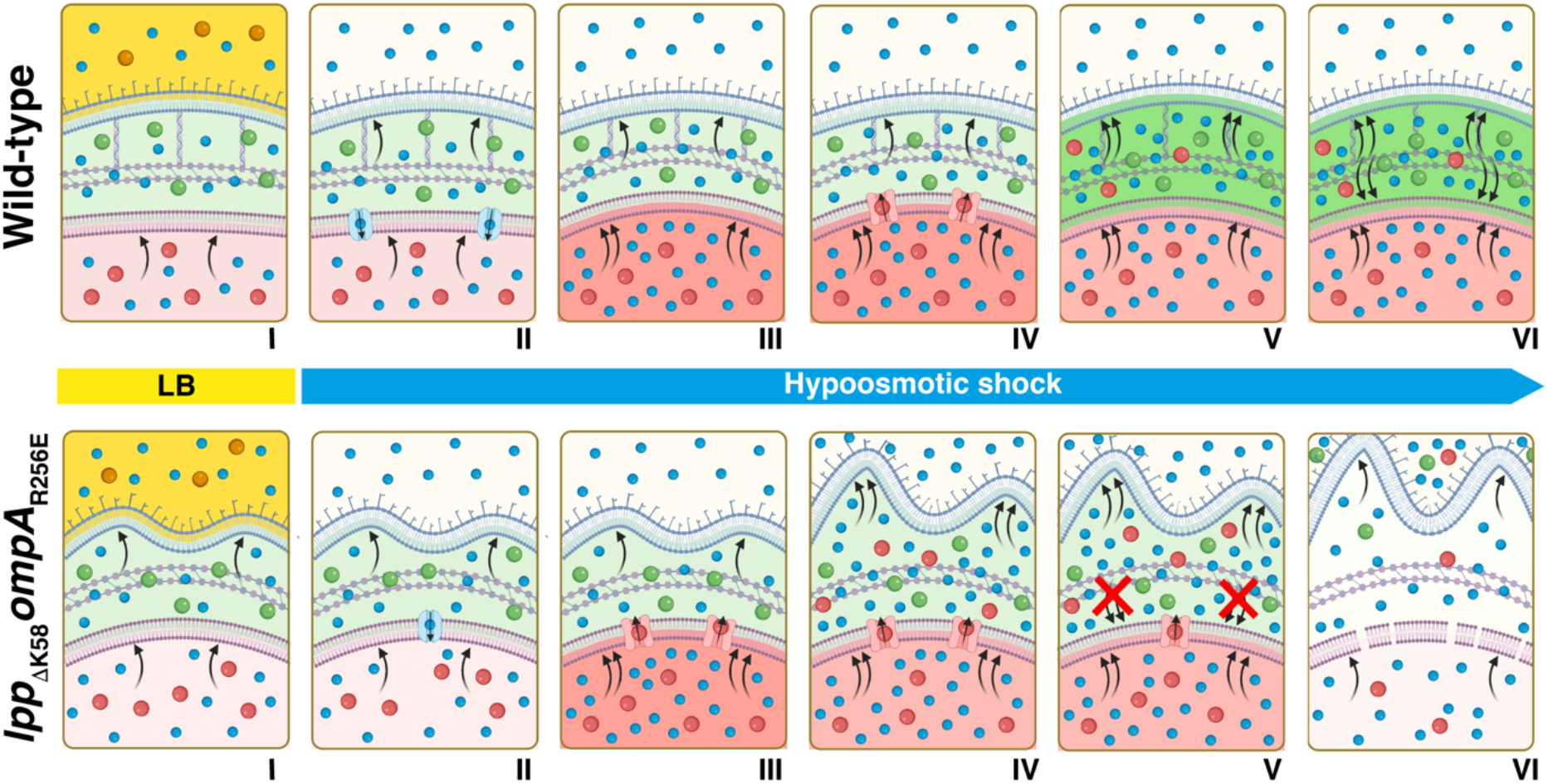
Attaching the outer membrane to the peptidoglycan layer is required for osmoresistance. Upon an osmotic down-shift, rapid water (blue circles) influx into the cytoplasm (step II), mediated by the aquaporin AqpZ, increases turgor. The resulting outward pressure on the cytoplasmic membrane (step III) activates the MSCs, which release osmotically active solutes (red circles) to the periplasm (step IV). In turn, the accumulation of solutes in the periplasm attracts more water molecules inside this compartment. In wild-type cells (upper panels), the outer membrane is attached to the peptidoglycan layer, which limits the expansion of periplasmic volume and allows periplasmic turgor (dark green) to build up (step V). An inward pressure is then applied on the cytoplasmic membrane, balancing cytoplasmic turgor and maintaining membrane integrity (step VI): these cells resist hypoosmotic shocks. In *lpp*_ΔK58_*ompA*_R256E_ cells (lower panels), the influx of water in the periplasm exerts an outward pressure on the outer membrane (step IV), which pushes the membrane apart from the cell body. This increases the periplasm volume, leading to the formation of a large bulge and the release of soluble content to the environment (step V), preventing the build-up of periplasmic turgor. No inward pressure is applied on the cytoplasmic membrane, which becomes destabilized and releases cytosolic components to the periplasm; thus, these cells die (step VI).

### Effect of strengthening the disconnected outer membrane

This model poses that the connection between the outer membrane and the peptidoglycan layer is crucial because it allows the build-up of periplasmic turgor (*28*), which counterbalances the outward pressure on the cytoplasmic membrane, protecting its integrity and preventing cell death. In this view, attaching the outer membrane to the peptidoglycan layer allows the periplasm to function as a counter pressure chamber. Thus, we postulated that consolidating the outer membrane in the *lpp*_ΔK58_*ompA*_R256E_ mutant should compensate for the lack of attachment and allow the build-up of the periplasmic turgor, at least to a certain extent. If our postulate is true, reinforcing the outer membrane might increase the resistance of the *lpp*_ΔK58_*ompA*_R256E_ mutant to hypoosmotic shocks. Excitingly, this is what we observed. To strengthen the outer membrane, we incubated cells with MgCl_2_ or CaCl_2_ (1 mM); these cations bridge lipopolysaccharide (LPS) molecules by preventing the repulsion of their phosphate groups (*29*). Moreover, 1 mM MgCl_2_ corresponds to ∼3 mOsm, which is far below cellular osmolality (hundreds to thousands mOsm (*30*)). Remarkably, we found that adding Mg^2+^ or Ca^2+^ rescues the resistance of the *lpp*_ΔK58_*ompA*_R256E_ mutant to osmotic down-shifts (Fig. 3A) and reduces the release of intracellular proteins to the medium (Fig. 3B, fig. S5), thus supporting our model. The addition of an isotonic concentration of NaCl or NiCl had no impact, indicating that the rescue does not result from the increased osmolality of the MgCl_2_ or CaCl_2_ solutions but from the ability of Mg^2+^ or Ca^2+^ to coordinate the phosphate groups of the LPS molecules. Analysis at the single-cell level corroborates these results: 83.8%±11.4% (n=3132 cells) of the Mg^2+^-strengthened cells survived treatment, maintaining cytoplasmic mCherry inside the cytoplasm (88.9%±7.9%) with an unperturbed localization pattern of GlpT (Fig. 3C, fig. S7, movies S5–6). The outer membrane of surviving cells forms bulges (in 83.6% of cells) in which msfTq2^C70V^ accumulate (Fig. 3B). However, despite bulge formation, the cytoplasmic membrane is not perturbed and mCherry is not transferred to the periplasm; the cells do not die (fig. S7B).

**Fig. 3.**
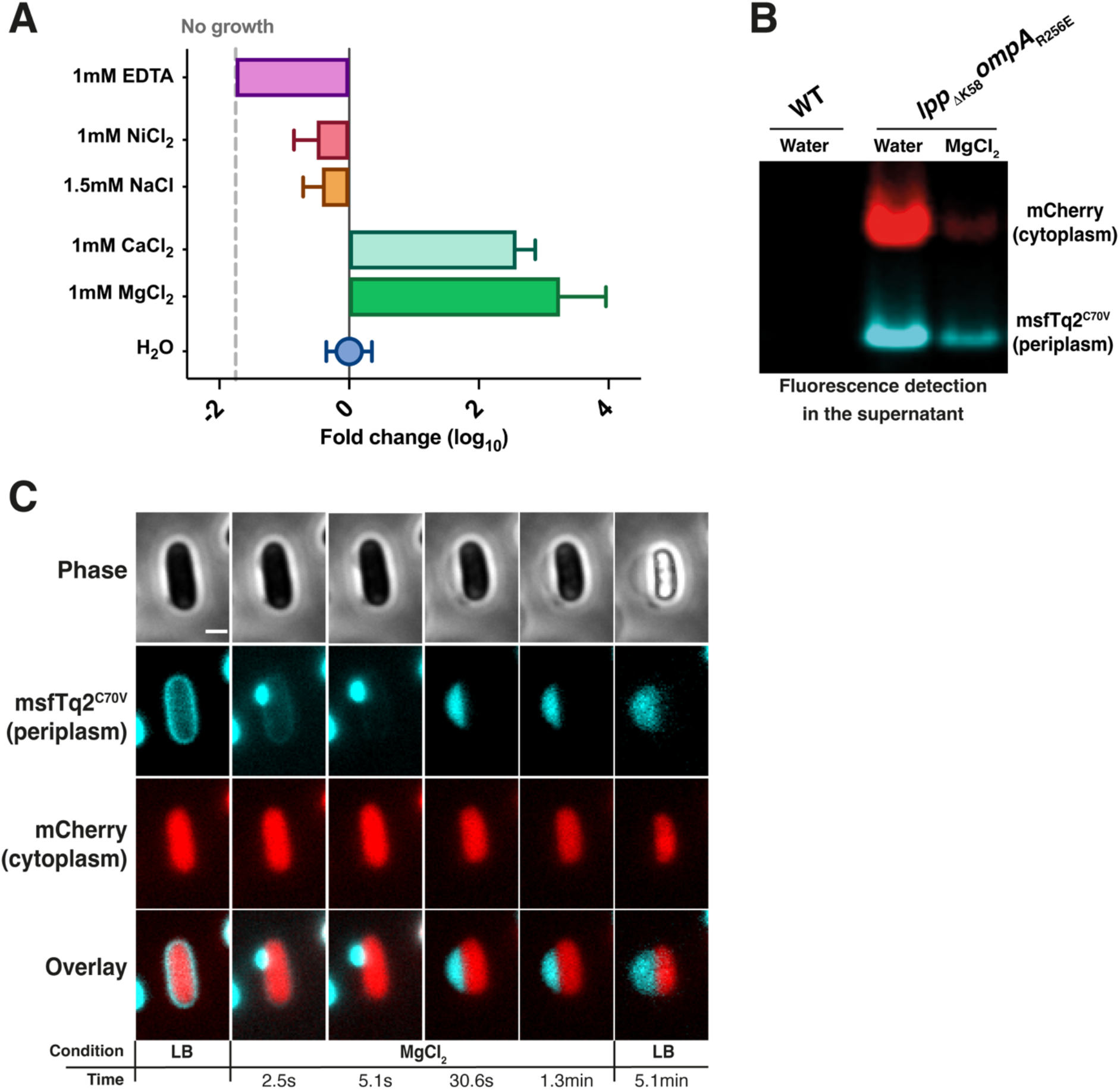
Strengthening the outer membrane increases the resistance of *lpp*_ΔK58_*ompA*_R256E_ cells to hypoosmotic shocks. (**A**) Incubation with MgCl_2_ and CaCl_2_, but not with NaCl or NiCl_2_, reduces the sensitivity of *lpp*_ΔK58_*ompA*_R256E_ cells to hypoosmotic shocks —Mg^2+^ and Ca^2+^, but not Na^+^ or Ni^2+^, bridge LPS molecules, preventing their repulsion. Incubation with the salt chelator ethylenediaminetetraacetic acid (EDTA) further increases osmosensitivity. *lpp*_ΔK58_*ompA*_R256E_ cells were resuspended on hypotonic medium supplemented with the indicated salts or chelators and serially diluted. Dilutions were spotted on an LB agar plate. (**B**) Mg^2+^ addition decreases the release of periplasmic msfTq2^C70V^ and cytoplasmic mCherry to the supernatant when *lpp*_ΔK58_*ompA*_R256E_ cells are exposed to a hypoosmotic down-shift. Cell supernatants after hypoosmotic shock were loaded on a native gel and imaged by fluorescence. WT: wild type. (**C**) The impact of Mg^2+^ on the ability of cells with an impaired outer membrane–peptidoglycan connection to withstand hypoosmotic shifts was analyzed using microfluidics. *lpp*_ΔK58_*ompA*_R256E_ cells expressing msfTq2^C70V^ (periplasm) and mCherry (cytoplasm) were shifted from LB to a MgCl_2_-supplemented hypotonic medium for 5 min before being shifted back to LB. Snapshots were taken at the indicated times. Mg^2+^ addition does not prevent the formation of a bulge in which msfTq2^C70V^ accumulates; this bulge keeps the periplasmic content inside the periplasm and mCherry inside the cytoplasm. Most cells (83.8%±11.4%) respond to the osmotic upshift, indicating cell survival. Scale bar: 2 μm. Images were acquired every 2.55 s.

### Effect of water flux across the cytoplasmic membrane

To further test our model, we sought to manipulate the water flux across the cytoplasmic membrane. Indeed, in our model, the triggering event of the deadly cascade observed in mutant cells is the rapid influx of water into the cytoplasm upon an osmotic down-shift. Previous research has shown that the rate of osmotic down-shock determines the survival probability of MSC mutants (*31*). Thus, our model predicts that preventing water from rapidly entering the cytoplasm should have a protective effect, limiting the opening of MSCs and the release of solutes to the periplasm, which should reduce the sensitivity of the *lpp*_ΔK58_*ompA*_R256E_ mutant to osmotic down-shifts. To decrease the water flux, we deleted *aqpZ*, the gene encoding the only aquaporin expressed by *E. coli*. Aquaporins greatly enhance cytoplasmic water influx; in their absence, water only slowly diffuses through the cytoplasmic membrane (*32*). Excitingly, we found that *aqpZ* deletion substantially reduces the sensitivity of the *lpp*_ΔK58_*ompA*_R256E_ mutant to hypoosmotic shocks (Fig. 4A), while overexpression of *aqpZ*, but not of an inactive mutant (AqpZ_R189S_)(*33*), further sensitizes cells (Fig. 4A). In line with these results, a lower level of intracellular proteins was released by the *lpp*_ΔK58_*ompA*_R256E_Δ*aqpZ* mutant upon water shock (Fig. 4D, fig. S5). Microfluidics provided insights into this behavior: we observed that *aqpZ* deletion in *lpp*_ΔK58_*ompA*_R256E_ cells delays the transfer of cytoplasmic components from the cytoplasm to the periplasm after water shock (mean or average time: ∼66.2 s instead of ∼30.1 s; Fig. 4B, fig. S8A–D, movies S7, S8, S9). Moreover, the addition of MgCl_2_ to *lpp*_ΔK58_*ompA*_R256E_Δ*aqpZ* cells drastically reduced (by 38.8%±18%) bulge formation (Fig. 4C, fig. S8A–B, E–F, movie S10) and intracellular protein release (Fig. 4D, fig. S4).

**Fig. 4.**
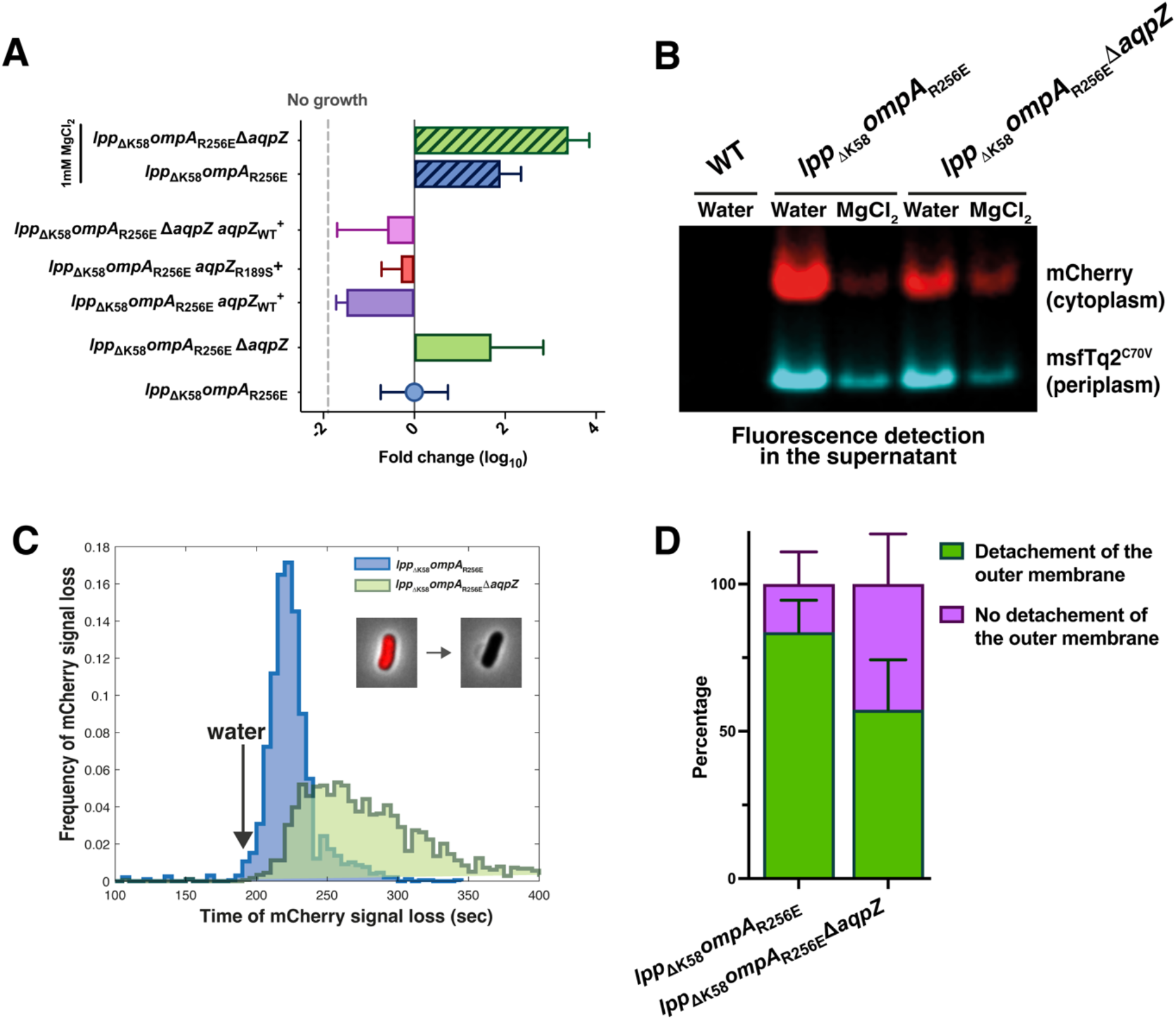
Decreasing water influx into the cytoplasm reduces the sensitivity of cells with a disconnected outer membrane to hypoosmotic shock. (**A**) Deleting the gene encoding the aquaporin AqpZ reduces the sensitivity of *lpp*_ΔK58_*ompA*_R256E_ cells to hypoosmotic shock, while AqpZ overexpression further sensitizes these cells. Expression of a catalytic mutant (AqpZ_R189S_) has no impact. Adding MgCl_2_ further reduces the osmosensitivity of *lpp*_ΔK58_*ompA*_R256E_Δ*aqpZ* cells compared with *lpp*_ΔK58_*ompA*_R256E_. Cells were resuspended in hypotonic medium, serially diluted, and spotted on an LB agar plate. (**B**) Deleting *aqpZ* decreases the release of periplasmic msfTq2^C70V^ and cytoplasmic mCherry to the supernatant when *lpp*_ΔK58_*ompA*_R256E_ cells are exposed to a hypoosmotic down-shift. Addition of Mg^2+^ further reduces the release. Cell supernatants after hypoosmotic shock were loaded on a native gel and imaged using fluorescence. WT: wild type. (**C**) Deleting *aqpZ* delays the release of mCherry to the cell exterior by 53 s (mean loss of 43.7 s and 96.7 s for *lpp*_ΔK58_*ompA*_R256E_ [n=1977 cells] and *lpp*_ΔK58_*ompA*_R256E_Δ*aqpZ* [n=1788 cells], respectively). Loss of the mCherry signal during a hypoosmotic down-shift was monitored and timed using microfluidics. (**D**) Deleting *aqpZ* substantially decreases (by 38.8%) the number of *lpp*_ΔK58_*ompA*_R256E_ cells in which the outer membrane detaches from the cell body when shifted to a Mg^2+^-containing hypotonic solution. *lpp*_ΔK58_*ompA*_R256E_ (n=2784 cells) and *lpp*_ΔK58_*ompA*_R256E_Δ*aqpZ* (n=3313 cells) cells expressing periplasmic msfTq2^C70V^ and cytoplasmic mCherry were analyzed using microfluidics.

## DISCUSSION

It is textbook knowledge that the peptidoglycan cell wall protects bacterial cells from bursting due to internal turgor pressure. This view, in which the peptidoglycan layer confers resistance against osmotic shocks by caging the cytoplasmic membrane, is mostly influenced by analogies with plant cells and concepts inferred for Gram-positive bacteria (*34*), in which a thick peptidoglycan layer is the only layer surrounding the cytoplasmic membrane. Here, we have shown that this model does not apply to Gram-negative bacteria. Instead, our results support a model in which the cell envelope as a whole protects these organisms from differences in osmotic pressure between the interior and exterior of the cell. In this view, the cell envelope functions as a pressurized chamber, in which pressure build-up counterbalances cytoplasmic turgor, thereby maintaining cytoplasmic membrane integrity. Our model also suggests that periplasmic turgor is required to allow MSC closure following sudden osmotic down-shifts and proposes that periplasmic turgor is the driving force for the formation of outer membrane vesicles (*13*). The importance of the outer membrane– peptidoglycan connection in maintaining osmoprotection in *Pseudomonas aeruginosa* (fig. S9, movies S11–12) supports the idea that our model applies to all diderm bacteria.

Our work also highlights the role of outer membrane–peptidoglycan connecters. Lpp, the numerically most abundant protein in *E. coli*, is known to be important for outer membrane integrity —the outer membranes of cells lacking lpp form blebs and detach from the cell body (*11*)— and for dictating the size of the envelope (*7, 35*). The importance of the tethering function of OmpA has been less clear, although the functional relevance of the interaction between its periplasmic domain and the peptidoglycan layer has been demonstrated (*36*). Here, we reveal a novel, perhaps the most crucial, function of these two proteins, which is to constrain the size of the envelope compartment to allow the build-up of periplasmic turgor. Interestingly, *E. coli*, like many other bacteria (*37*), expresses Pal, an outer membrane lipoprotein with peptidoglycan-binding properties (*38*). We were unable to create a Δ*pal* mutant in *lpp*_ΔK58_*ompA*_R256E_ cells, suggesting that a minimal level of outer membrane–peptidoglycan connection is required for growth and survival, opening a field for future research.

A long-standing mystery is the evolutionary transition between the cell envelopes of diderm and monoderm bacteria (*39*). In a recent study, it was proposed that monoderm bacteria evolved from diderm bacteria and that the triggering factor was the loss of the outer membrane–peptidoglycan connecters (*10*). Within the framework of our model, it is tempting to speculate that maintaining a detached outer membrane was sufficient to provide an osmotically protective environment in certain organisms until a thicker peptidoglycan cell wall progressively evolved.

By discovering that the outer membrane–peptidoglycan connection is required for Gram-negative bacteria to withstand cytoplasmic turgor, our work reveals the previously unsuspected role of the outer membrane in osmoresistance and provides an answer to the question of why the outer membrane is essential. Our study also sheds new light on crucial molecular mechanisms that allow bacteria to adapt and survive in often hostile environments, an understanding of which is required for the development of innovative antimicrobial strategies.

## ACKNOWLEDGMENTS

We thank Abel Garcia-Pino and Ariel Talavera Perez for assistance with cryo-EM experiments and Wendy Le Mouellic for helping with exploratory experiments. We are indebted to members of the lab for helpful suggestions and discussions and to Géraldine Laloux for providing comments on the manuscript. We are indebted to Tanneke den Blauwenn for providing plasmids to Romé Voulhoux for *P. aeruginosa* strains. We thank the staff at the VIB-VUB Bio Electron Cryogenic Microscopy facility for assistance in data collection. This work was funded by Fonds de la Recherche Scientifique (FNRS) grant agreements (WELBIO-CR-2015A-03, WELBIO-CR-2019C-03), the Flanders Research Foundation Hercules grant (G0H5916N), the Flanders Research Foundation PhD fellowship program, and VIB. We thank Kaneka Corporation for financial support.

## Author contributions

M.D. and J.-F.C. wrote the paper; M.D., S.-H.C., and J.-F.C. conceived the project and designed the study; M.D., S.-H.C., S.G., A.J., and A.D. performed the experiments; M.D., S.- H.C., S.G., A.J., H.R., and J.-F.C. analyzed the data.

## Competing interests

The authors declare no competing interests.

## Data and materials availability

All data are available in the main text or the supplementary materials or upon request.

## SUPPLEMENTARY MATERIALS

Materials and Methods

## Notes

### Competing Interest Statement

The authors have declared no competing interest.

