## Supplemental Material and methods for "The outer membrane and peptidoglycan layer form a single mechanical device balancing turgor"

**The PDF file includes:**

Materials and Methods

### Materials and Methods

#### Bacterial strains, plasmids, and culture

The bacterial strains and plasmids used in this study are listed in tables S1 and S2. We obtained gene containing point mutation of OmpA$_{R256E}$ by allelic replacement using λRed recombination (*40*, *41*). For subsequent transfer of these mutations in other strains, we used phage P1 transduction by first introducing a kanamycin resistance cassette originating from the Keïo collection (*42*) in non-coding regions located close to the genes of interest (i.e., between *fumD* and *pikF* and between *matP* and *ompA* for Lpp and OmpA, respectively). *torD::kan* (from the Keïo collection) was introduced in JLE126. We performed subsequent co-transfer of *torD::kan* and *ompA::ompA*$_{R256E}$ in other strain backgrounds using phage P1 transduction. When required, we removed the *kan* gene using site-specific recombination induced by expression of the Flp recombinase from pCP20 plasmid (*40*). All primers are listed in table S3.

pASKA-aqpZ$_{R189S}$ was obtained by site-directed mutagenesis of the ASKA plasmid collection containing the *aqpZ* gene (*43*). We cloned the genes coding for cytoplasmic mCherry into pTHV037-$_{ss}$DsbA-sfTq2$^{C70V}$ using Gibson assembly (*44*).

We applied Gibson assembly to clone the gene coding for cytoplasmic mCherry in Ptrc-msfGFP-4GS-GlpT-Bla-pSC101 (*45*). We also cloned the gene coding for lpoB-mCherry (*44*) in Ptrc-msfGFP-4GS-GlpT-Bla-pSC101 using Gibson assembly. Bacteria were grown in LB-Miller formulation (10 g/L NaCl). When appropriate, we used supplements at the following concentrations: ampicillin (Ap) 100 μg/mL or 200 μg/mL, chloramphenicol (Cm) 20 μg/mL, kanamycin (Kn) 50 μ g/mL, streptomycin (St) 20 μ g/mL, isopropyl β-d-1-thiogalactopyranoside (IPTG) 15 μM or 100 μM, arabinose (ara) 0.2%.

When appropriate, cells were cultivated with 100 μM of IPTG for the induction of *aqpZ* and *aqpZ* mutated genes or with 10 μM for the induction of fluorescent proteins after the optical density at 600 nm (OD$_{600nm}$) reached 0.1.

#### Hypoosmotic down-shift survival assay

Exponential phase cultures of bacterial resuspension were spun down, and extensive removal of the supernatant was performed. The pellet was resuspended to twice its original volume in the appropriate buffer (e.g., pure water, 1 mM $MgCl_2$, phosphate-buffered saline) for 5 min and then serially diluted in the same buffer and plated on an LB agar plate. We measured the survival rate by counting the colony number at the highest-fold dilution and rounding off at 0.5. We calculated the mean and standard deviation of the survival colony number for at least three replicates. For each graph, the survival rate of the reference strain was set to 0 (represented by a circle). We determined the ratio of cell survival of a tested strain or condition by dividing the corresponding number by the number of the reference strain or condition. We displayed the results as histograms using GraphPad Prism software. Negative ratios indicate a higher mortality of the corresponding strain or condition compared with the reference, while positive ratios indicate a higher survival rate. We calculated the survival limit rate from the survival rate of the reference strain or condition (i.e., 0) minus the mean of this reference strain or condition.

#### Microfluidics

We imaged cells on a Nikon Ti2-E fully motorized inverted epifluorescence microscope equipped with a CFI Plan Apochromat λ DM 100x1.45/0.13 mm Ph3 oil objective (Nikon), a

Sola SEII FISH illuminator (Lumencor), and a Prime95B camera (Photometrics). We controlled the multi-dimensional image acquisition using NIS-Ar software (Nikon). Bacterial strains were cultivated until they reached exponential phase at 37°C in LB medium supplemented with the appropriate inducer when needed and were then loaded onto B04A microfluidic perfusion plates (CellASIC). We performed medium modification using the ONIX microfluidics platform (CellASIC). Cells were first flushed in LB medium (containing IPTG when appropriate) for 3 min and then shifted to the tested buffer (e.g., pure water, 10 mM $MgCl_2$, LB) for 5 min before being resuspended in LB. Cytoplasmic condensation upon LB resuspension was used to determine survival (*23*). We performed medium exchange at 2 psi for each buffer and strain. Images were acquired every 500 ms.

**Oufti analysis**

*Extraction of cellular characteristics.* We obtained cell contours from phase contrast images using the open-source software package Oufti (*46*). Fluorescent signals were added to the obtained cell outlines after background subtraction within Oufti. Morphological features (e.g., cell length, width, area, and volume) and fluorescence information (e.g., signal intensities) were determined by summing the dimensions of each individual segment of the cell mesh identified by Oufti and stored in the Oufti cellList data structure. See https://oufti.org/ for more details.

*Determination of signal loss during time-lapse experiments.* To determine the timing of fluorescent signal loss, we monitored fluorescence concentrations over time using the obtained cellLists in MATLAB. To account for photobleaching, which led to a gradual reduction in fluorescence intensities over time, we monitored fluorescence signals between consecutive frames and looked for sudden decreases in the fluorescence signal, indicative of cells losing their corresponding perisolic or cytosolic content. The timing of the signal loss was taken as the first frame in which the fluorescence concentration was 1.4 times lower than the average of the two previous frames (corresponding to a signal reduction of >28.5%). Cells with low fluorescence concentrations in the first frame (0.1 arbitrary units (AU)/$\mu m^3$) were excluded from the analysis, as reductions in the fluorescence signal over time could no longer be reliably detected.

**Cryo-EM**

For cryo-EM analysis, we prepared sample grids by spotting 3 µL of bacteria ($OD_{600nm}$=0.5) on glow-discharged Lacey carbon films on copper 300 mesh (Agar Scientific). The grids were manually back-blotted (3–4 s) and flash-frozen in liquid ethane using a CP3 cryoplunger (Gatan). We collected images on a 300-kV CRYO ARM™ 300 (JEM-Z300FSC) field emission cryo-EM (JEOL) equipped with a K3 Summit direct electron detector (Gatan) at the VIB-VUB Bio Electron Cryogenic Microscopy facility in Brussels, Belgium. The detector was used in integrating mode with a 5-s exposure time at a nominal magnification of 5000x, corresponding to a pixel size of 1.021 nm.

**Native gel**

Exponential phase cultures (500 µL) of strains SMDeg977, SMDeg978, and SMDeg1307 induced with 10 µM IPTG were subjected to a hypoosmotic down-shift with 1 mL of water or 1 mM $MgCl_2$ (described previously). Supernatant supplemented with 2 µg bovine serum albumin for loading control was concentrated on a Vivaspin column at 30-fold, mixed (1:1) with native Tris-glycine sample buffer (Invitrogen), and loaded onto Novex Tris-glycine gels (Invitrogen). We visualized the fluorescence bands using a Hamersham ImageQuant 800 camera with Cy5 (mCherry) and Cy2 (msfTq2ox) fluorescence filters.

**Protein release assay**

*Analysis of the pellet and supernatant of water-shock cells.* We grew 2.5 mL of wild-type, *lpp*$_{\Delta K58}$*ompA*$_{R256E}$, and *lpp*$_{\Delta K58}$*ompA*$_{R256E}$ *ΔaqpZ* cells in LB-Miller medium and harvested 1 mL of cells at an $OD_{600}$ of ~0.5 by centrifugation at 9391g for 1 min. A 0.5-mL sample of each of the three supernatants (Fraction A) was kept for analysis, while the rest was discarded, and the cell pellet was used for the water shock. Two similar samples of *lpp*$_{\Delta K58}$*ompA*$_{R256E}$ and *lpp*$_{\Delta K58}$*ompA*$_{R256E}$ *ΔaqpZ* cells were also prepared for the magnesium ion-containing water shock (1 mM $MgCl_2$). We added 2 mL of water to the pellets of wild-type, *lpp*$_{\Delta K58}$*ompA*$_{R256E}$, and *lpp*$_{\Delta K58}$*ompA*$_{R256E}$ Δ*aqpZ* cells and 2 mL of 1 mM $MgCl_2$ to the pellets of *lpp*$_{\Delta K58}$*ompA*$_{R256E}$ and *lpp*$_{\Delta K58}$*ompA*$_{R256E}$ *ΔaqpZ* cells. The pellets were mildly resuspended, maintained for 5 min at room temperature, and centrifuged at 9391g for 1 min. A 1-mL sample of each of the five supernatants (Fraction B) was kept for analysis, while the rest was discarded, and the cell pellets (Fraction C) were kept for analysis. For Fractions A and B, 60% trichloroacetic acetic acid (TCA; final concentration: 10%) was added to precipitate proteins. For Fraction C, 200 µL of 10% TCA was added. All treated samples in Fractions A, B, and C were kept on ice for 30 min and centrifuged at 16,863g for 3 min. The obtained precipitates were treated with 400 µL of acetone and centrifuged at 16,863g for 5 min. Sodium dodecyl-sulfate polyacrylamide gel electrophoresis (SDS-PAGE) sample buffer (50 mM Tris-HCl pH 7.5, 0.15 M NaCl, 10% glycerol, 0.002% bromophenol blue, 1% SDS) was added to the precipitates (200, 200, and 400 µL for the samples from Fractions A, B, and C, respectively). We loaded 10 µL from the 13 samples onto the SDS-PAGE gel to analyze the outer membrane, inner membrane, periplasmic, and cytoplasmic proteins using Western blot.

*Western blot and antibodies.* Using a semi-dry electroblotting system, we transferred proteins from the gels onto nitrocellulose membranes (Millipore) for Western blot analyses, which were performed in the standard format. The signals were detected by chemiluminescence from the reaction of horseradish peroxidase with luminol. Rabbit anti-DsbA and anti-DsbD antibodies have been previously used by our group (*47*, *48*). Because the anti-DsbA antibody had been affinity-purified and diluted with 50 mM Tris-HCl buffer, pH 7.5, we used a 1:50 dilution, and we used anti-DsbD antibody at a dilution of 1:20,000. We used rabbit anti-OmpC (A64454, EpiGentek) at a dilution of 1:10,000, rabbit anti-TrxA (T0803, Sigma) at 1:10,000, rabbit anti-RpsB (CSB-PA08644A0Rb, Cusabio) at 1:20,000, and mouse anti-EF-Tu (HM6010, Hycult Biotech) at 1:10,000.

**Software**

Cartoons and schematics were generated using BioRender and Adobe Illustrator.

**Wild-type and transposon mutants of *Pseudomonas aeruginosa* UCBPP-PA14**

OprI is a homolog of Lpp in *P. aeruginosa* PA01 (*49*). Earlier work reported that Oprl was absent in PA14 because of an early stop, but it was later shown that Oprl is present and is expressed in PA14 (*50*, *51*). However, in the PA14 transposon mutant library, an OprI mutant is missing. Thus, we used a mutant of a peptidoglycan-crosslinking enzyme (l,d transpeptidase) as a defective peptidoglycan-crosslinking mutant of OprI. In *P. aeruginosa* PA14, PA27180 (annotated as erfK [ldtA]) appears to be the only gene homologous to Ec*ldtA, ldtB,* and *ldtC* (*52*). Therefore, we used the PA27180 gene transposon insertion mutant.
